# Compartment-Specific Lysosomal Heterogeneity in Microglia Is Regulated by Adaptor Protein Complex-4

**DOI:** 10.64898/2026.09.17.752408

**Authors:** Elizabeth A. Somodji, Swetha Gowrishankar

**Affiliations:** Department of Anatomy and Cell Biology, College of Medicine, University of Illinois Chicago, Chicago, IL 60612, USA; Department of Biological Sciences, College of Liberal Arts and Sciences, University of Illinois Chicago, Chicago, IL 60607, USA

## Abstract

Microglia rely on lysosomes to clear phagocytosed material and orchestrate immune responses, but the mechanisms governing glial lysosome diversity and function remain largely undefined. Using high-resolution, high-throughput 3D imaging of mouse brain tissue, we find that glial lysosomes are heterogeneous in size and protease content depending on subcellular location. CD68+ and LAMP1+ vesicles were more numerous in glial processes than cell bodies; process LAMP1+ vesicles carried less protease than cell-body vesicles, but CD68+ protease content did not differ between compartments, revealing at least two lysosomal subpopulations within processes. Loss of Adaptor Protein complex 4 (AP-4) selectively remodels this landscape: it increases CD68+ vesicle number and CD68 enrichment, particularly in processes, without changing protease content, and disrupts the normal cell-body-versus-process pattern in protease- and CD68-enrichment. AP-4 loss also lowers protease content in cell-body LAMP1+ vesicles, mirroring a neuronal phenotype. These results provide the first evidence of glial lysosome heterogeneity and implicate AP-4 in shaping microglial lysosome composition and inflammatory function.

## Introduction

Lysosome dysfunction is closely linked to several neurodegenerative diseases. Of the CNS cell types, microglia, the resident phagocytic cells of the brain with high degradative demands, are perhaps most sensitive to lysosome dysregulation. Indeed, further linking glial lysosome dysfunction with neurodegenerative disease, multiple recent studies have revealed that distinct lysosomal insults can trigger an epigenetic program that is also observed in disease states, including in Alzheimer’s Disease (AD) (Balak et al., 2026; Chavan and Bhattacharjee, 2025; Tejwani et al., 2026). Several lines of evidence also support strong links between immune activation and changes to lysosome biogenesis and functioning. This includes both network-level changes to lysosomal pathways on immune activation as well as changes to specific endo-lysosomal proteins when immune-related proteins such as TREM2 are perturbed (Borst et al., 2021; Filipello et al., 2023; Iyer et al., 2022; Wang et al., 2024). One cellular feature that has drawn attention in many other cell types, which may contribute to the organelle’s versatility in microglia by equipping it to process diverse substrates based on the functional demand, is lysosomal heterogeneity. Heterogeneity in signaling, degradative capacity, and enrichment of certain components (Akter et al., 2023; Bond et al., 2025; Narendra et al., 2020; Settembre and Perera, 2024) has been well documented in cultured cells alongside a growing appreciation for its relevance to cell physiology and disease pathology, when disrupted (Chen and Gutierrez, 2025; Paumier and Gowrishankar, 2024; Settembre and Perera, 2024; Somodji and Gowrishankar, 2026). More recently, lysosome heterogeneity has also been described based on subcellular positioning (Bussi and Gutierrez, 2024; Gowrishankar and Ferguson, 2016; Paumier and Gowrishankar, 2024; Pu et al., 2016). Despite this, the mechanisms by which this operates in the natural 3D environment, particularly in professional degraders such as microglia, remain unstudied. To this end, we have undertaken an in-depth, high-resolution study of microglial endo-lysosomes within intact mouse brain tissue. By combining high-speed, high-resolution imaging with in-depth 3D image analysis and mutant studies, we interrogate whether spatial and compositional glial lysosome heterogeneity exists under wildtype conditions and if so, how this is disrupted by loss of the Adaptor Protein-4 complex (AP-4). Our prior studies of AP-4 KO human neurons and a mouse model have revealed that loss of the complex exerts distinct effects on different neuronal lysosome populations, altering lysosome composition in soma lysosomes versus potentially altering transport properties of axonal lysosomes (Majumder et al., 2022). We hypothesized that microglia, with their high degradative capacity, morphological complexity (Barrella et al., 2025; De Biase et al., 2017; Lawson et al., 1990; Vidal-Itriago et al., 2022) that includes formation and retraction of highly dynamic processes, are likely to have compositionally, functionally distinct lysosomes in these different subcellular locations, which in turn would be differentially regulated by AP-4. We found that CD68+ vesicles, while smaller and more numerous in processes than in the main glial body, retained comparable protease content, a departure from the canonical high in soma, low in processes gradient we and others have reported in neurons (Cheng et al., 2018; Gowrishankar et al., 2015; Yap et al., 2018), and even in non-polarized cell lines (Johnson et al., 2016). This could suggest that CD68+ vesicles are a specialized subtype of lysosomes present in microglia and other macrophages. We also found that AP-4 loss drives striking changes to process-specific CD68+ vesicles, increasing not just their abundance, but also their CD68 and cathepsin L enrichment. Interestingly, LAMP1+ vesicles exhibited a neuron like gradient in protease enrichment, and AP-4 loss decreased cathepsin L content in lysosomes within the glial body. To our knowledge, this is the first study to characterize microglial CD68+ lysosomes as heterogeneous based on their subcellular position within microglia in situ, and to show that this heterogeneity is a distinct axis from the well-established position-dependent heterogeneity of LAMP1+ organelles seen in other cell types, and now also in glia. The divergent effects of AP-4 loss on CD68+ and LAMP1+ vesicles suggest that distinct routes of sorting/biogenesis exist in microglia, with CD68+ vesicles not acquiring cathepsin L through AP-4 dependent biosynthetic delivery. The AP-4 dependent changes driven specifically in processes, including CD68 protein enrichment, vesicle abundance, cathepsin L increase which has been mechanistically tied to TNF-α induction in other systems (Xu et al., 2018), support a model wherein these distal CD68+ compartments are poised for a proinflammatory/secretory role, and that AP-4 dysfunction (already linked to altered plaque-microglia engagement in an AD model) may act partly by driving this process-localized inflammatory lysosomal phenotype. More broadly, these findings position subcellular lysosomal heterogeneity as a previously overlooked variable in microglial biology, one that may determine how genetic and disease-associated perturbations translate into region-specific degradative or inflammatory outputs within a single cell.

## Results

### CD68+ vesicles are protease-rich and differ in abundance based on their cellular position within microglia

CD68, a type 1 transmembrane protein exclusively labels microglial lysosomes in the brain (Bornemann et al., 2001; Holness et al., 1993; Rabinowitz and Gordon, 1991), and is widely used as an identifier for activated, phagocytic glia (Ayata et al., 2018; Bornemann et al., 2001; Hendrickx et al., 2017; Hopperton et al., 2018; Nicoll et al., 2006). Changes in CD68 expression and number of CD68+ glia have been linked to both AD and physiological ageing (Hopperton et al., 2018; Tsering et al., 2025; Wong et al., 2005). More recently, glial subpopulations have even been identified within an individual brain based on CD68 content, suggesting that CD68 enrichment contributes to microglial heterogeneity (Ayata et al., 2018; Hendrickx et al., 2017; Wharton et al., 2015; Wong et al., 2005). However, the nature of CD68+ lysosomes, including their content and distribution within an individual glia, are not known. Likewise, our prior studies revealed that not only does AP-4 regulate neuronal lysosome composition (Majumder et al., 2022), its loss lead to altered glial recruitment to amyloid plaques in an AD model (Orlowski et al., 2024), a feature linked to the cell’s autophagy-lysosomal pathways (Choi et al., 2023). Given this, we stained and imaged CD68 along with Iba1 (which labels the entire microglial structure) in the brains of both WT and AP-4 KO mice and analyzed the high-resolution images of glia we acquired from within the hippocampus CA1 region using the 3D analysis software, IMARIS. After utilizing the Iba1 labeling as a “mask” to delineate each individual glia (Video 1) and demarcating glial processes from their “main body”, we analyzed CD68+ vesicles within these subcellular domains in 3D (Video 2, 3). Firstly, we found that even within individual WT glia (Fig. 1A, Video 3), the CD68+ vesicles exhibit heterogeneity as a function of their position within the cell. We observed that there were far more, but smaller CD68+ vesicles within processes as compared to the main body (Fig. 1B, C). However, these vesicles tended to be equally enriched in CD68 as those in the main body (Fig. 1D). Thus, it appears that while the abundance of CD68+ vesicles is indeed influenced by cellular positioning, certain other properties do not change.

**Figure 1.**
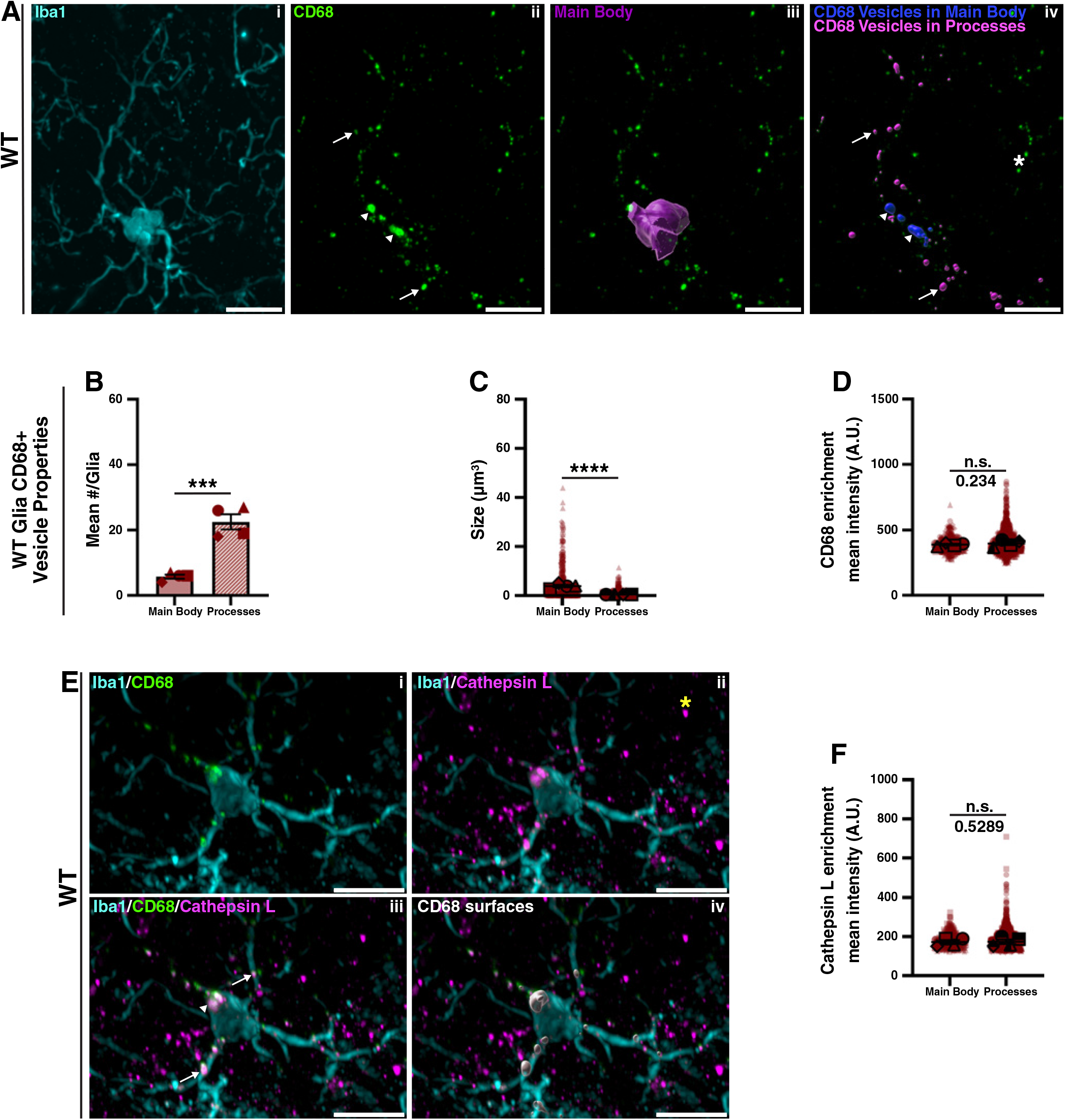
CD68+ vesicles in WT microglia are enriched in cathepsins, and exhibit heterogeneity based on cellular positioning. (A) Representative high-resolution images depicting a single WT microglia. Iba1 [(i) glial body; cyan] was used to identify microglial cell bodies, while CD68 [(ii) CD68+ vesicles; green] was used to identify microglial endo-lysosomes in the main body (white arrowheads) and processes (white arrows). (iii) Image showing “main body” (purple surface) of microglia and CD68+ vesicles (green). (iv) Image showing CD68+ vesicles in 3D or as rendered “surfaces” within the main body (arrowheads; blue surfaces) and in processes (arrows; magenta surfaces). Note, CD68+ vesicles outside this individual microglia’s body are visible in green (white asterisk) and not included in the glia’s CD68+ vesicle pool (blue or magenta). Bar, 10 µm. (B) Quantification of mean number of CD68+ vesicles per microglia within the main body and processes of WT microglia in the hippocampus CA1 region of 8-month-old male AP-4 WT animals (main body - solid bar; processes-striped bar). Each shape (circle, square, triangle, and diamond) represents data from an individual animal. Mean ± SEM, N = 4 animals, n = 66 glia. *** p< 0.001, unpaired *t*-test. (C) Superplot depicting CD68+ vesicle size, as measured by volume (µm³), of all individual CD68+ vesicles from the main body and processes from microglia of the individual WT animals (circle, square, triangle, and diamond), as well as the mean CD68+ vesicle volume of each animal (larger symbol with bold outline) ± SEM. N = 4 animals, n = 379 CD68+ vesicles in the main body and n = 1,456 CD68+ vesicles in the processes. **** p<0.0001. Shown are *p*-values from a Linear Mixed Effects (LME) model, (D) Superplot depicting CD68 enrichment, as measured by mean intensity (arbitrary intensity units), of all individual CD68+ vesicles from the main body and processes from microglia of the individual WT animals (circle, square, triangle, and diamond), as well as the mean CD68 intensity of each animal (larger symbol with bold outline) ± SEM. N = 4 animals, n = 379 CD68+ vesicles in the main body and n = 1,456 CD68+ vesicles in processes. ns= not significant. Data were analyzed using an LME model. (E) High-resolution images highlighting (i) a microglia (Iba1; cyan) in the WT hippocampus CA1 region co-stained with CD68 (green) labeling endo-lysosomes within the glia; (ii) highlighting the cathepsin L (magenta), which labels vesicles within the glia (Iba1; cyan) and also neighboring neurons (yellow asterisk). (iii) Merged image that depicts Iba1, CD68 and cathepsin L highlighting vesicles within the main body (white arrowheads) and processes (white arrows) of glia. (iv) Image of Iba1, CD68 and cathepsin L, where the isolated CD68 vesicles from within the individual glia (identified by its Iba1 mask) are rendered in 3D or as white “surfaces”. The cathepsin L content from within these surfaces are specifically analyzed to capture properties relating to the individual microglia’s vesicles. Bar, 10 µm. (F) Superplot depicting cathepsin L enrichment, as measured by mean intensity of cathepsin L (arbitrary intensity units), of all individual CD68+ vesicles in the main body and processes from microglia of the individual WT animals (circle, square, triangle, and diamond), as well as the mean cathepsin L intensity of each animal (larger symbol with bold outline) ± SEM. N = 4 animals, n = 379 CD68+ vesicles in the main body and n = 1,456 CD68+ vesicles in the processes. ns= not significant. Data were analyzed using an LME model.

In certain cell lines as well as highly polarized neurons, other key properties of LAMP1+ vesicles such as their acidification and protease content have been shown to differ as a function of their position (Cheng et al., 2018; Gowrishankar et al., 2015; Gowrishankar and Ferguson, 2016; Hollenbeck, 1993; Johnson et al., 2016; Kulkarni and Maday, 2018; Majumder et al., 2022; Yap et al., 2018). In neurons, lysosomes within the neuronal cell body are highly enriched in proteases, while the more distal processes are relatively protease-poor (Cheng et al., 2018; Gowrishankar et al., 2015; Yap et al., 2018). We therefore compared the protease content within WT glia CD68+ vesicles in the main body versus processes that had been stained for cathepsins L and B (Fig. 1E, F, Fig. S1 A, B; Video 4). Intriguingly, we found no difference in cathepsin L or B enrichment within CD68+ lysosomes based on their position within microglia (Fig. 1F and Fig. S1 B). Thus, CD68+ vesicles within processes could have the same degradative potential as those within the main body, a departure from observations of lysosomes in neurons (Cheng et al., 2018; Gowrishankar et al., 2017, 2015; Yap et al., 2018). The presence of cathepsins in these vesicles at levels similar to the main body would suggest that these CD68+ vesicles at the very least, are phagolysosomes/phagosomes that have fused with endo-lysosomes, if in fact they are phagosomal rather than endosomal in origin.

### Loss of AP-4 alters CD68+ vesicle properties and increases cathepsin L in glial lysosomes in processes

Lysosomes across different regions are heterogeneous not just in composition, but also in their sensitivity to distinct perturbations (Bussi and Gutierrez, 2024; Gowrishankar and Ferguson, 2016; Paumier and Gowrishankar, 2024; Pu et al., 2016). Indeed, we previously found that loss of AP-4 in neurons, affected the composition and proteolytic function of neuronal soma lysosomes, while it appeared to alter the motility and clearance of lysosomes from axonal processes (Majumder et al., 2022). Not only is the AP-4 complex a regulator of neuronal lysosome composition and function, but we also found that its loss altered microglial recruitment to amyloid plaques in a mouse model of Alzheimer’s Disease (Orlowski et al., 2024). This, along with prior findings that optimal proteostasis is required for appropriate plaque-glia engagement (Choi et al., 2023), suggest that the AP-4 complex could also regulate microglial lysosomal properties. To address this, we examined how loss of AP-4 impacted glial CD68+ vesicle properties, utilizing the tissue from AP-4 KO (Fig. S2 A, Video 5) that we imaged along with their respective sex-matched WT littermates. We find that loss of AP-4 function altered multiple aspects of microglial CD68+ vesicles when compared to WT. Firstly, there was a striking increase in CD68+ vesicle abundance in AP-4 KO glia overall (Fig. 2A, B, Fig. S2 B-D, Video 6, Video 7). When we parsed this vesicle abundance based on their sub-cellular position (Fig. 2C-E), we found that while the abundance of CD68+ vesicles within main body of AP-4 KO didn’t differ from those of the WT (Fig. 2C), their abundance within AP-4 KO microglial processes was significantly increased (Fig. 2D). We found no differences between AP-4 WT and KO, in terms of the number of processes associated with individual glia (Fig. S2 E), nor did the glial volumes differ (Fig. S2 F), suggesting that this is a true increase in CD68+ vesicle abundance rather than an increase in number or size/complexity of processes. This is an intriguing similarity to the increased lysosome abundance within axonal processes (Majumder et al., 2022), though we did not see any dystrophies in glial processes or an aggregation of CD68+ lysosomes within areas of them. Given the links between CD68 and inflammation (Chistiakov et al., 2017; Hopperton et al., 2018; Wong et al., 2005), this increase in CD68+ vesicles upon loss of AP-4 function, could imply a heightened activation state of these glia. At the individual vesicle level, loss of AP-4 did not change the size of CD68+ vesicles in either compartment when compared to WT glia (Fig. 2F, G); however, it did lead to differences in the size of CD68+ vesicles between the main body and processes within AP-4 KO glia themselves (Fig. 2H), similar to CD68+ vesicles within WT glia (Fig. 1C). Loss of AP-4 complex function also increased CD68 enrichment on individual vesicles compared to WT, regardless of cellular positioning (Fig. 2I, J). Another interesting alteration of glial lysosomes due to AP-4 loss, is that it disrupts cell body versus processes lysosomal properties: unlike in the WT, AP-4 loss led to higher CD68 enrichment on the vesicles in the glial processes when compared to vesicles in the AP-4 KO main body (Fig. 2K). Thus, in addition to a striking increase in abundance of CD68+ vesicles within processes, loss of AP-4 introduces heterogeneity in CD68 enrichment between cell body and process lysosomes, fundamentally altering lysosomal diversity within the cell. Given links of CD68 to phagocytic activity, oxidative stress and inflammatory response (Hopperton et al., 2018; Tsering et al., 2025; Wong et al., 2005), these stark changes in CD68 vesicle properties could indicate that these glia are more activated and/or phagocytic in nature.

**Figure 2.**
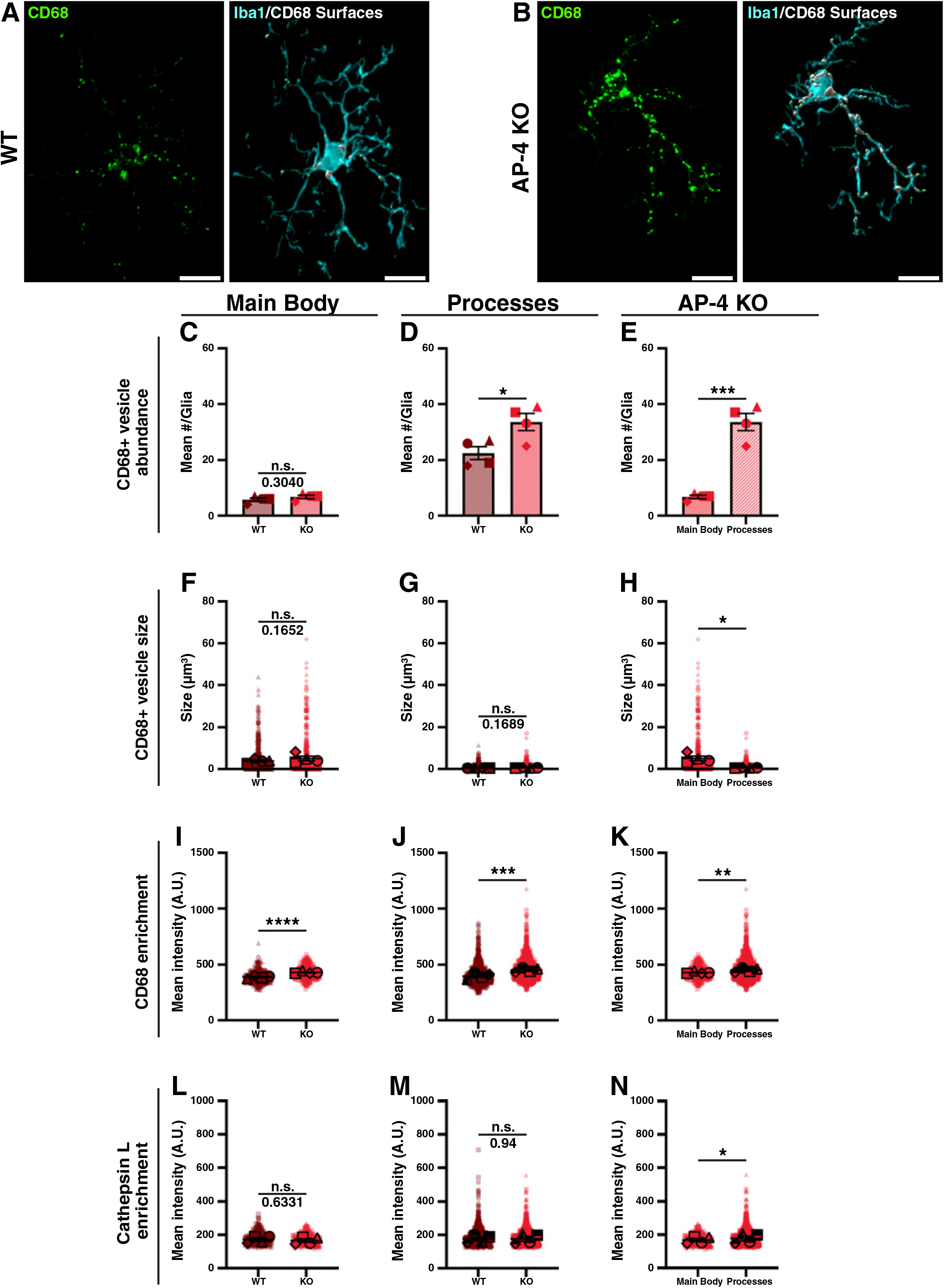
Loss of AP-4 alters CD68+ vesicle properties and content specifically in glial processes. (A, B) Representative high-resolution “mask” images of microglia from AP-4 WT(A) and KO (B) hippocampus CA1 region, stained to label microglial cell bodies (Iba1; cyan) and microglial endo-lysosomal vesicles (CD68; green), wherein use of Iba1 surface as a mask, isolates CD68+ vesicles from within the individual glia alone (green signal, white surface renderings). Bar, 10 µm. (C) Quantification of mean number of CD68+ vesicles per microglia within the main bodies of AP-4 WT and KO microglia in the hippocampus CA1 region of 8-month-old male AP-4 WT and KO animals (WT – brown; KO – red). Each shape (circle, square, triangle, and diamond) represents data from an individual animal. Mean ± SEM, N = 4 animals, n = 66 WT glia, n = 57 AP-4 KO glia. Ns = not significant, unpaired *t*-test. (D) Quantification of mean number of CD68+ vesicles per microglia within the processes of AP-4 WT and KO microglia in the hippocampus CA1 region of 8-month-old male AP-4 WT and KO animals (WT – brown; KO – red). Each shape (circle, square, triangle, and diamond) represents data from an individual animal. Mean ± SEM, N = 4 animals, n = 66 WT glia, n = 57 AP-4 KO glia. * p<0.05, unpaired *t*-test. (E) Quantification of mean number of CD68+ vesicles per microglia within the main body and processes of AP-4 KO microglia in the hippocampus CA1 region of 8-month-old male AP-4 KO animals (main body-solid bar; processes-striped bar). Each shape (circle, square, triangle, and diamond) represents data from an individual animal. Mean ± SEM, N = 4 animals, n = 57 glia. *** p< 0.001, unpaired *t*-test. (F) Superplot depicting CD68+ vesicle size, as measured by volume (µm³), of all individual CD68+ vesicles from the main bodies from microglia of the individual AP-4 WT and KO animals (circle, square, triangle, and diamond), as well as the mean CD68+ vesicle volume of each animal (larger symbol with bold outline) ± SEM. N = 4 animals, n = 379 CD68+ vesicles in WT glial main bodies; n = 396 CD68+ vesicles in AP-4 KO glial main bodies. ns = not significant. Data were analyzed using an LME model. (G) Superplot depicting CD68+ vesicle size, as measured by volume (µm³), of all individual CD68+ vesicles from the processes from microglia of the individual AP-4 WT and KO animals (circle, square, triangle, and diamond), as well as the mean CD68+ vesicle volume of each animal (larger symbol with bold outline) ± SEM. N = 4 animals, n = 1,456 CD68+ vesicles in WT processes; n = 1,924 CD68+ vesicles in AP-4 KO processes. ns = not significant. Data was analyzed using an LME model. (H) Superplot depicting CD68+ vesicle size, as measured by volume (µm³), of all individual CD68+ vesicles from the main body and processes from microglia of the individual AP-4 KO animals (circle, square, triangle, and diamond), as well as the mean CD68+ vesicle volume of each animal (larger symbol with bold outline) ± SEM. N = 4 animals, n = 396 CD68+ vesicles in the main body; n = 1,924 CD68+ vesicles in the processes. * p<0.05. Shown are *p*-values from an LME model.(I) Superplot depicting CD68 enrichment, as measured by mean CD68 intensity (arbitrary intensity units), of all individual CD68+ vesicles from the main bodies of microglia of the individual AP-4 WT and KO animals (circle, square, triangle, and diamond), as well as the mean CD68 intensity of each animal (larger symbol with bold outline) ± SEM. N = 4 animals, n = 379 CD68+ vesicles in WT glial main bodies; n = 396 CD68+ vesicles in AP-4 KO glial main bodies. **** p<0.0001. Shown are *p*-values from an LME model. (J) Superplot depicting CD68 enrichment, as measured by mean intensity (arbitrary intensity units), of all individual CD68+ vesicles from the processes of microglia of the individual AP-4 WT and KO animals (circle, square, triangle, and diamond), as well as the mean CD68 intensity of each animal (larger symbol with bold outline) ± SEM. N = 4 animals, n = 1,456 CD68+ vesicles in WT processes; n = 1,924 CD68+ vesicles in AP-4 KO processes. *** p<0.001. Shown are *p*-values from an LME model. (K) Superplot depicting CD68 enrichment, as measured by mean intensity (arbitrary intensity units), of all individual CD68+ vesicles from the main body and processes from microglia of the individual AP-4 KO animals (circle, square, triangle, and diamond), as well as the mean CD68 intensity of each animal (larger symbol with bold outline) ± SEM. N = 4 animals, n = 396 CD68+ vesicles in the main body; n = 1,924 CD68+ vesicles in the processes. ** p<0.01. Shown are *p*-values from an LME model. (L) Superplot depicting cathepsin L enrichment, as measured by mean intensity of cathepsin L (arbitrary intensity units), of all individual CD68+ vesicles from the main bodies of microglia from individual AP-4 WT and KO animals (circle, square, triangle, and diamond), as well as the mean cathepsin L intensity of each animal (larger symbol with bold outline) ± SEM. N = 4 animals, n = 379 CD68+ vesicles in WT main bodies and n = 396 CD68+ vesicles in AP-4 KO main bodies. ns = not significant. Data was analyzed using an LME model. (M) Superplot depicting cathepsin L enrichment, as measured by mean intensity of cathepsin L (arbitrary intensity units), of all individual CD68+vesicles from the processes of microglia from individual AP-4 WT and KO animals (circle, square, triangle, and diamond), as well as the mean cathepsin L intensity of each animal (larger symbol with bold outline) ± SEM. N = 4 animals, n = 1,456 CD68+ vesicles in WT processes and n = 1,924 CD68+ vesicles in AP-4 KO processes. ns = not significant. Data was analyzed using an LME model. (N) Superplot depicting cathepsin L enrichment, as measured by mean intensity of cathepsin L (arbitrary intensity units), of all individual CD68+ vesicles from the main body and processes of microglia of the individual AP-4 KO animals (circle, square, triangle, and diamond), as well as the mean cathepsin L intensity of each animal (larger symbol with bold outline) ± SEM. N = 4 animals, n = 396 CD68+ vesicles in the main body; n = 1,924 CD68+ vesicles in the processes. * p<0.05. Shown are *p*-values from an LME model.

A comparison of protease content of CD68+ vesicles between WT and AP-4 KO glia revealed that there was no appreciable differences between them, both in the main body and processes (Fig. 2L, M). This is different from our findings in neurons, where AP-4 loss in neurons also results in a decrease of cathepsin L in lysosomes, albeit characterized there based on their LAMP1 positivity (Majumder et al., 2022). The cathepsin B content in CD68+ vesicles was also not altered in either vesicle population of AP-4 KO glia when compared to WT (Fig. S2 G, H). Interestingly, we found that cathepsin L was enriched in AP-4 KO CD68+ vesicles in the processes compared to those in AP-4 KO main body (Fig. 2N), which again contrasts with our findings in the WT glia (Fig. 1F). This protease enrichment is likely more specific rather than a global change to protease content as cathepsin B content was not altered between the main body and processes CD68+ vesicles in AP-4 KO glia (Fig. S2 I).

Our results thus reveal a previously unappreciated heterogeneity of CD68+ lysosomes where, based on their cellular position, they differ in abundance and response to perturbations.

The similar protease content in CD68+ vesicles in WT glial processes compared to the main body vesicles contrasts with the findings in both neurons (Cheng et al., 2018; Gowrishankar et al., 2015; Yap et al., 2018). and cultured cell lines (Johnson et al., 2016), where the potentially equivalent axonal, dendritic lysosomes or more distal lysosomes in non-polarized cultured cells have relatively lower protease content compared to the cell body/more perinuclear lysosomes. None of the studied examples, however, were positive for CD68, suggesting that these CD68+ compartments may be a specialized subtype of lysosomes seen in microglia and macrophages. While the higher protease content could mean increased degradative capacity, cathepsins B and L expression have also been linked to inflammation (Lowry and Klegeris, 2018; Somodji and Gowrishankar, 2026). Studies in cell lines and mice implicate cathepsin B in maturation and processing of IL-1β (Jiang et al., 2025; Terada et al., 2010), and increased expression of cathepsin L in LPS-stimulated BV2 cells was linked to expression of proinflammatory TNF-α (Xu et al., 2018). In both cases, lysosomal release of these proteases has been correlated with proinflammatory responses (Liu et al., 2008; Ni et al., 2019). It is tempting to speculate that the CD68+ vesicles in glial processes (and thus more distal ones in the cell) are involved in inflammatory responses and are prone to secretion. Consistent with this possibility, loss of AP-4 complex function, which is associated with disturbed neuro-proteostasis, increases the abundance, CD68+ enrichment, and cathepsin L content of these CD68+ vesicles within glial processes. The increased abundance and differences in content raise questions about the biogenesis of these organelles as well as their engagement (through fusion and fission) with other endo-membranes. Likewise, future studies involving co-staining with ILs and other inflammation markers could help further functionally delineate this lysosomal population.

### LAMP1+ vesicles show position-dependent differences in content and properties within microglia, with AP-4 loss driving reduced cathepsin enrichment in the organelles

Lysosomal-associated membrane protein 1 (LAMP1) is an abundant glycoprotein on the surface of both late endosomes and degradative lysosomes (Ballabio and Bonifacino, 2020; Chen et al., 1985; Lewis et al., 1985; Saftig and Klumperman, 2009), known to be enriched on compositionally, functionally heterogenous organelles, in both non-polarized cultured cell lines (Johnson et al., 2016) and polarized primary cells such neurons (Ballabio and Bonifacino, 2020; Bussi and Gutierrez, 2024; Cheng et al., 2018; Gowrishankar et al., 2017, 2015; Yap et al., 2018). We thus examined how distinct LAMP1+ vesicles were distributed across compartments within a glia and how loss of AP-4 function affected them. We stained LAMP1, cathepsin L and Iba1 (to label the entire microglial structure) in the brains of both WT and AP-4 KO mice, and imaged and analyzed them as we did previously for CD68 vesicles (Figs 1, 2). Comparisons of distribution and properties of LAMP1 + vesicles in distinct locations within glia revealed that while they shared some features with CD68+ vesicles, there were other clear differences. In WT glia, as with CD68+ vesicles, LAMP1+ vesicles were far more numerous within the processes when compared to the main body (Fig. 3A, B, Fig. S3 A, Video 8). LAMP1+ vesicles in processes were also smaller but exhibited less enrichment of LAMP1 protein on individual vesicles when compared to those in the main body (Fig. 3C, Fig. S3 B). Interestingly, we observed LAMP1+ vesicles in the main body exhibited more enrichment of cathepsin L when compared to the population in processes (Fig. 3D), which is consistent with observations in neurons, but different with CD68+ vesicles, where organelles in processes were just as enriched with cathepsins as those in the main body (Fig. 1F). Thus, heterogeneity based on cellular position of LAMP1+ vesicles is distinct from that of CD68+ vesicles. LAMP1+ vesicles in AP-4 KO glia exhibited similar properties to WT glia, in that the LAMP1+ vesicles in their processes were more abundant (Fig. 3E, F, Fig. S3 C, Video 9), smaller in size (Fig. 3G), less enriched in LAMP1 protein (Fig. S3 D), and relatively cathepsin L-poor (Fig. 3H). LAMP1+ vesicles in AP-4 KO glia not only exhibit the same position-dependent heterogeneity as WT glia, in the above-described properties, they do not differ from WT glia in their abundance or size in either main body or processes (Fig. 3I-L, Fig. S3 E). This again is a stark contrast to the increased number of CD68+ vesicles in AP-4 KO glia. We also did not observe a change in LAMP1 enrichment between AP-4 WT and KO glia (Fig. S3 F, G). However, AP-4 loss did decrease cathepsin L enrichment within LAMP1+ vesicles of the main body specifically, when compared to WT glia (Fig. 3M, N), a point of phenotypic convergence with AP-4 KO neurons (Majumder et al., 2022). Thus, LAMP1+ vesicles differ from CD68+ vesicles in their position-based heterogeneity as well as how they are altered due to loss of AP-4. This would suggest that at the very least, LAMP1+ and CD68+ vesicles do not completely overlap in microglia. This is further supported by higher number of LAMP1+ vesicles in processes (Fig.3B) as compared to CD68+ vesicles (Fig.1B). The presence of a relatively protease-deficient population of LAMP1+ vesicles within processes of glia of either genotype as observed in distal axons of neurons in the brain tissue (Gowrishankar et al., 2015) would suggest these are less mature, non-degradative compartments that could progressively mature as they move further into the main body. AP-4 loss affecting cathepsin L enrichment within LAMP1 vesicles in the main body would suggest an enzyme sorting defect/delivery as was observed in neurons (Majumder et al., 2022). However, AP-4 KO glia do not lack cathepsin enrichment within CD68+ vesicles, including in processes. We thus propose that the CD68+ vesicles do not acquire their cathepsin content by the same biosynthetic delivery route as with LAMP1+ compartments in the main body of glia, but from fusion with other endo-lysosomal intermediates and/or mechanisms that are integrated with immune activation-dependent cathepsin synthesis. Together, our results establish subcellular position and molecular identity (CD68 versus LAMP1) as two independent axes of lysosomal heterogeneity in microglia, and show that a single trafficking perturbation, namely AP-4 loss, remodels these two compartments in distinct and compartment-specific ways rather than uniformly disrupting glial lysosome function (Fig. 4). Future studies that extend this analysis in a disease context, such as Alzheimer’s Disease and in physiological ageing, would help establish whether the process-selective remodeling of CD68+ lysosomes we observe in AP-4 KO mice is a general feature of microglial activation states relevant to neurodegeneration and/or ageing, or a phenotype specific to loss of AP-4 function.

**Figure 3.**
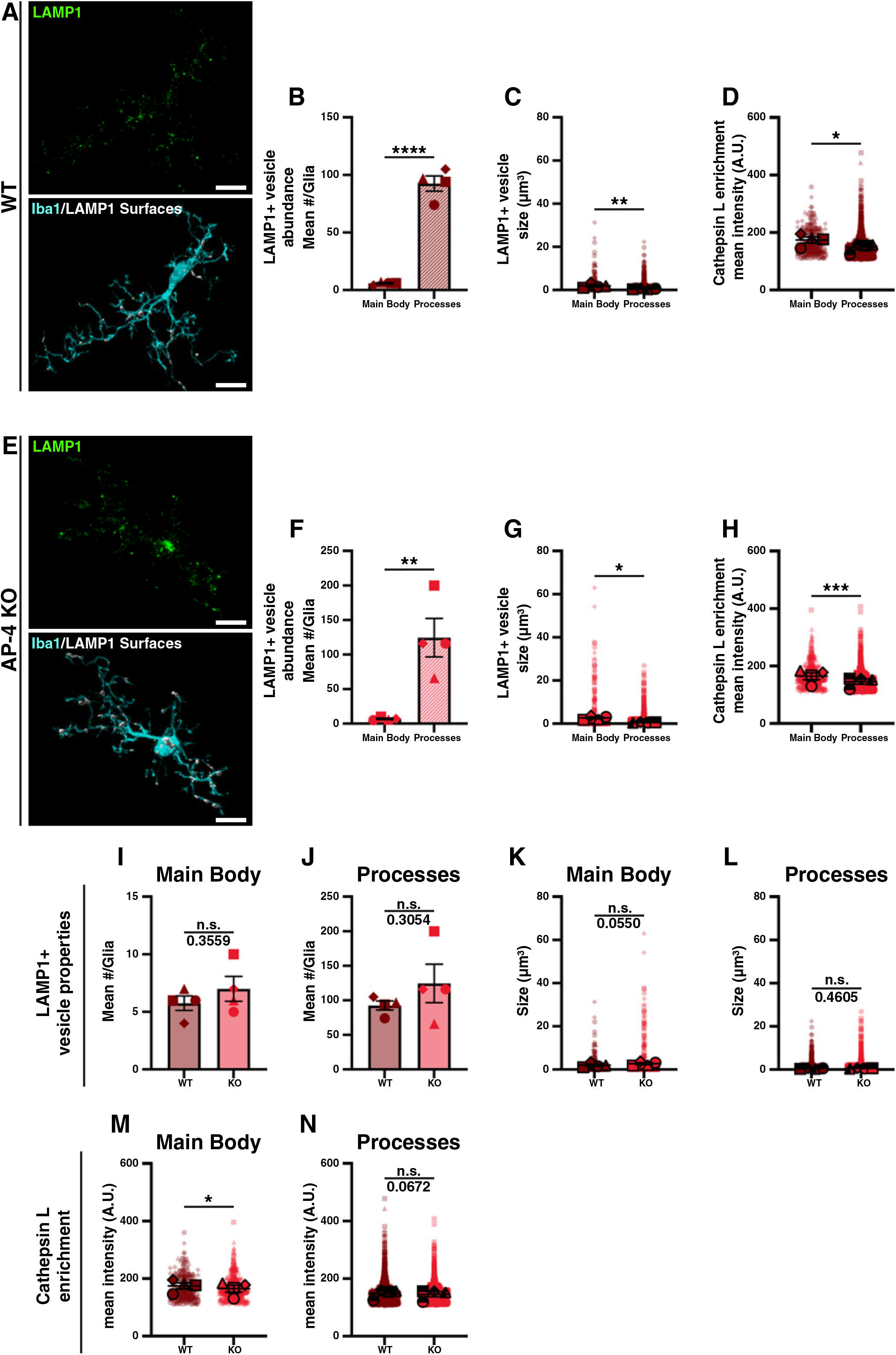
AP-4 loss differentially affects LAMP1+ vesicle properties as compared to CD68+ vesicles. (A) Representative high-resolution mask image of a microglia from hippocampus CA1 region of an AP-4 WT mouse stained for the microglial cell body (Iba1; cyan) and endo-lysosomes (LAMP1; green), wherein use of Iba1 surface as a mask, isolates LAMP1+ vesicles from within the individual glia alone (green signal, white surface renderings). Bar, 10 µm. (B) Quantification of mean number of LAMP1+ vesicles per microglia within the main body and processes of microglia in the hippocampus CA1 region of AP-4 WT animals (main body – solid bar; processes – striped bar). Each shape (circle, square, triangle, and diamond) represents data from an individual animal. Mean ± SEM, N = 4 animals, n = 42 glia. **** p<0.0001, unpaired *t*-test. (C) Superplot depicting LAMP1+ vesicle size, as measured by volume (µm³), of all individual LAMP1+ vesicles from the main body and processes of microglia from the individual WT animals (circle, square, triangle, and diamond), as well as the mean LAMP1+ vesicle volume of each animal (larger symbol with bold outline) ± SEM. N = 4 animals, n = 246 LAMP1+ vesicles in the main body; n = 3,727 LAMP1+ vesicles in the processes. ** p<0.01. Shown are *p*-values from an LME model. (D) Superplot depicting cathepsin L enrichment, as measured by mean intensity of cathepsin L (arbitrary intensity units), of all individual LAMP1+ vesicles from the main body and processes of microglia of the individual WT animals (circle, square, triangle, and diamond) as well as the mean cathepsin L intensity of each animal (larger symbol with bold outline) ± SEM. N = 4 animals, n = 246 LAMP1+ vesicles in the main body; n = 3,727 LAMP1+ vesicles in the processes. * p<0.05. Shown are *p*-values from an LME model. (E) Representative high-resolution mask image of a microglia from hippocampus CA1 region of an AP-4 KO mouse stained for the microglial cell body (Iba1; cyan) and endo-lysosomes (LAMP1; green), wherein use of Iba1 surface as a mask, isolates LAMP1+ vesicles from within the individual glia alone (green signal, white surface renderings). Bar, 10 µm. (F) Quantification of mean number of LAMP1+ vesicles per microglia within the main body and processes of KO microglia in the hippocampus CA1 region of 8-month-old male AP-4 KO animals (main body – solid bar; processes – striped bar). Each shape (circle, square, triangle, and diamond) represents data from an individual animal. Mean ± SEM, N = 4 animals, n = 57 glia. ** p<0.01, unpaired *t*-test. (G) Superplot depicting LAMP1+ vesicle size, as measured by volume (µm³), of all individual LAMP1+ vesicles from the main body and processes of microglia of the individual AP-4 KO animals (circle, square, triangle, and diamond), as well as the mean LAMP1+ vesicle volume of each animal (larger symbol with bold outline) ± SEM. N = 4 animals, n = 423 LAMP1+ vesicles in the main body; n = 7,625 LAMP1+ vesicles in the processes. * p<0.05. Shown are *p*-values from an LME model. (H) Superplot depicting cathepsin L enrichment, as measured by mean intensity of cathepsin L (arbitrary intensity units), of all individual LAMP1+ vesicles from main body and processes of microglia of the individual AP-4 KO animals (circle, square, triangle, and diamond) as well as the mean cathepsin L intensity of each animal (larger symbol with bold outline) ± SEM. N = 4 animals, n = 423 LAMP1+ vesicles in the main body; n = 7,625 LAMP1+ vesicles in the processes. *** p<0.001. Shown are *p*-values from an LME model. (I) Quantification of mean number of LAMP1+ vesicles per microglia within the main bodies of WT and KO microglia in the hippocampus CA1 region of 8-month-old male AP-4 WT and KO animals (WT – brown; KO – red). Each shape (circle, square, triangle, and diamond) represents data from an individual animal. Mean ± SEM, N = 4 animals, n = 42 WT glia, n = 57 AP-4 KO glia. ns = not significant, unpaired *t*-test. (J) Quantification of mean number of LAMP1+ vesicles per microglia within the processes of WT and KO microglia in the hippocampus CA1 region of 8-month-old male AP-4 WT and KO animals (WT – brown; KO – red). Each shape (circle, square, triangle, and diamond) represents data from an individual animal. Mean ± SEM, N = 4 animals, n = 42 WT glia, n = 57 AP-4 KO glia. ns = not significant, unpaired *t*-test. (K) Superplot depicting LAMP1+ vesicle size, as measured by volume (µm³), of all individual LAMP1+ vesicles from the main bodies of microglia of the individual AP-4 WT and KO animals (circle, square, triangle, and diamond), as well as the mean LAMP1+ vesicle volume of each animal (larger symbol with bold outline) ± SEM. N = 4 animals, n = 246 LAMP1+ vesicles in WT main bodies; n = 423 LAMP1+ vesicles in AP-4 KO main bodies. ns = not significant. Data was analyzed using an LME model. (L) Superplot depicting LAMP1+ vesicle size, as measured by volume (µm³), of all individual LAMP1+ vesicles from the processes of microglia of the individual AP-4 WT and KO animals (circle, square, triangle, and diamond), as well as the mean LAMP1+ vesicle volume of each animal (larger symbol with bold outline) ± SEM. N = 4 animals, n = 3,727 LAMP1+ vesicles in WT processes; n = 7,625 LAMP1+ vesicles in AP-4 KO processes. ns = not significant. Data was analyzed using an LME model. (M) Superplot depicting cathepsin L enrichment, as measured by mean intensity of cathepsin L (arbitrary intensity units), of all individual LAMP1+ vesicles from the main bodies of microglia of the individual AP-4 WT and KO animals (circle, square, triangle, and diamond) as well as the mean cathepsin L intensity of each animal (larger symbol with bold outline) ± SEM. N = 4 animals, n = 246 LAMP1+ vesicles in WT main bodies; n = 423 LAMP1+ vesicles in AP-4 KO main bodies. * p<0.05. Shown are *p*-values from an LME model. (N) Superplot depicting cathepsin L enrichment, as measured by mean intensity of cathepsin L (arbitrary intensity units), of all individual LAMP1+ vesicles from the processes of microglia of the individual AP-4 WT and KO animals (circle, square, triangle, and diamond) as well as the mean cathepsin L intensity of each animal (larger symbol with bold outline) ± SEM. N = 4 animals, n = 3,727 LAMP1+ vesicles in WT processes; n = 7,625 LAMP1+ vesicles in AP-4 KO processes. ns = not significant. Data were analyzed using an LME model.

**Figure 4.**
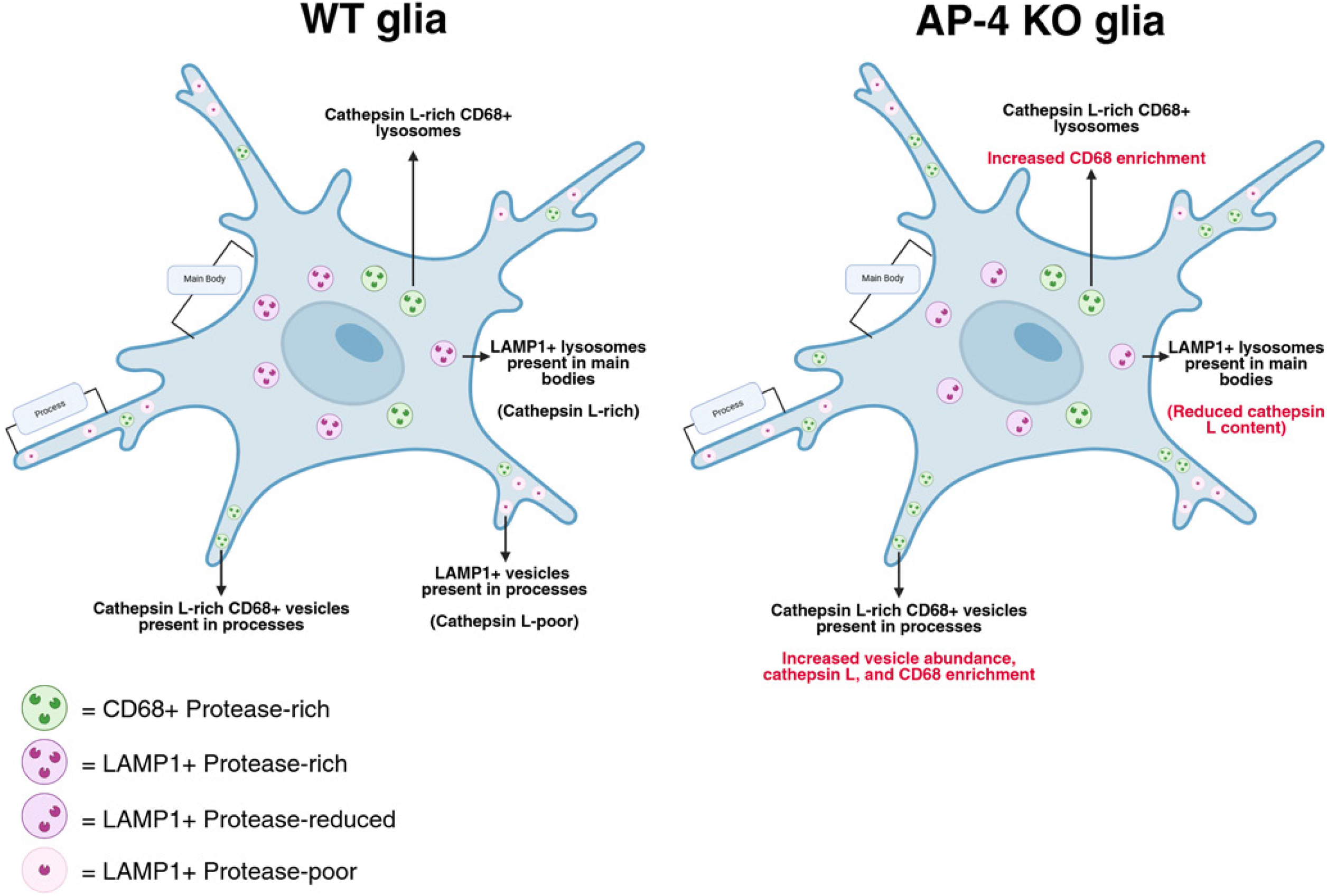
Compartment-specific lysosomal heterogeneity in microglia is regulated by Adaptor Protein Complex-4. Schematic of microglial lysosome subpopulations exhibiting two independent axes of lysosome heterogeneity in AP-4 WT glia and how this is altered in AP-4 KO glia, based on sub cellular position (main body and processes) and molecular identity (CD68 and LAMP1). In WT glia, both CD68+ and LAMP1+ vesicles appear to be smaller and more abundant in glial processes. However, the two subpopulations differ in their position-based heterogeneity based on vesicle content. CD68+ vesicles are cathepsin-rich regardless of their cellular position in glia, while LAMP1+ vesicles have differences in cathepsin L content similar to neurons, where vesicles in the processes are cathepsin L-poor while those in the main body are cathepsin L-rich. In AP-4 KO glia, CD68+ vesicles in the main body exhibit increase in CD68 enrichment on individual vesicles; however CD68+ vesicles within processes are more dramatically altered: there is an increase in their overall abundance, as well as an increase in cathepsin L-, and CD68-enrichment at the individual vesicle level. The changes to LAMP1 + vesicles due to AP-4 loss differ from those on CD68+ vesicles. Cathepsin L content within main body LAMP1+ vesicle is reduced in AP-4 KO glia, similar to what has been observed in AP-4 KO neurons.

## Methods

### Mice Strains

Animal procedures were approved by and carried out in accordance with guidelines established by the Office of Animal Care and Institutional Biosafety (OACIB). The C57BL/6N-Ap4ε1tm1b(KOMP)Wtsi (Davies et al., 2022; De Pace et al., 2018; Orlowski et al., 2024) was purchased from the European Mouse Mutant Archive (EMMA). Male and female heterozygous AP-4 mice were bred together as detailed before (Orlowski et al., 2024), to generate sex-matched AP-4 WT and KO littermates. 8-month-old male WT and KO littermates were used in this study. Progenies were genotyped by PCR in-house using the following primers: AP4E1-5arm-WTF(5′GCCTCTGTTTAGTTTGCGATG3′), AP4E1-Crit-WTR (5′CGTGCACAGACAGGTTTGAT3′), 5mut-R1(5′GAACTTCGGAATAGGAACTTCG3′).

### Transcardial Perfusion

Transcardial perfusion was performed to isolate tissue for immunohistochemistry experiments as described previously (Gowrishankar et al., 2017, 2015) 1xPBS was administered to anesthetized mice (Isoflurane) by transcardial perfusion. The isolated brains were then quickly dissected sagittally down the midline to procure hemibrains and one half fixed by immersion in 4% paraformaldehyde (PFA) overnight at 4°C while the other half was flash frozen in liquid nitrogen and stored at −80°C for future biochemical studies.

### Immunohistochemistry Studies

Hemibrains that were fixed in PFA overnight were washed three times in 1xPBS and sectioned coronally at 30 μm thickness using a vibratome (Leica VT1200S). Sections were collected serially into 8 wells of a 24-well plate containing 1xPBS with 0.05% sodium azide. Sections of the midbrain [corresponding to coronal sections 70–76 in the Allen Mouse Brain Atlas (atlas.brain-map.org)] were selected from the appropriate well and used for staining after matching them according to their depth. Staining experiments on the free-floating sections were performed as described previously (Gowrishankar et al., 2015). Antibodies used in this study are described in Table 1.

**Table 1.** Antibody Summary.

| Antibody | Source | Catalog Number | Dilution |
| --- | --- | --- | --- |
| Cathepsin B | R&D systems | AF965 | 1:200 |
| Cathepsin L | R&D systems | AF1515 | 1:200 |
| CD68 | Bio-Rad | MCA1957 | 1:200 |
| Iba1 | Wako | 019-19741 | 1:250 |
| LAMP1 | DSHB | 1D4B | 1:500 |

### Microscopy

16-bit 2-by-1 tile/stitched images of immunostained brain sections were acquired at 60X zoom using a 60x (NA 1.4) oil immersion objective on an Andor BC43 benchtop spinning disk confocal microscope (Andor Technology, Oxford Instruments) equipped with 405 nm, 488 nm, 561 nm, and 647 nm laser lines and an ZL41 cell 4.2 USB sCMOS camera. High-resolution Z-stacks were acquired with a step size of 0.1 µm using the Fusion software (Oxford Instruments). Tile images were saved as IMS files and stitched together using IMARIS Stitcher 10.2 (Oxford Instruments) and used for analysis.

### Quantitative Analysis (IMARIS)

#### CD68+ Analysis

30-μm coronal hemibrain sections obtained as described above, from 8-month-old AP-4 WT and KO male mice were stained for Iba1, CD68, and cathepsins L or B. Z-stacks of images that encompassed the Iba1 signal (typically 200-300 planes with 0.1 µm step size) and were within the same CA1 region of the hippocampus between AP-4 WT and KO mice, were analyzed using IMARIS 10.2 (Oxford Instruments). Iba1 intensity was corrected to reduce background noise using the *Normalize Layers* function and then the *Gaussian Filter* (width = 0.102 or 0.104) (Lind-Holm Mogensen et al., 2024). Glial cell bodies were isolated using the “*Surfaces”* feature with the Machine Learning Segmentation function on Iba1 signal where surfaces were then adjusted iteratively by manual training to ensure proper coverage of glia bodies and processes. The generated surfaces were then reviewed manually, so as to remove any glia/ surfaces that are cut off or incomplete within the acquired sections (glia that lie at the edges of Z stack captured and thus not fully encompassed within the stack). Following this, all the surfaces rendered and inspected were individually and manually edited (Nemes-Baran and DeSilva, 2021) to connect any segments that are segregated using “unify” and likewise separate objects that are independent but appear fused in the rendering, using “cut surface” function. Once the glial bodies are isolated, the CD68 channel/image was “masked” using the Iba1 surfaces, where signal from only within the completely rendered Iba1 surfaces is used to isolate CD68+ vesicles within these whole glia. CD68 surfaces were generated in the CD68 mask using automated surfaces (background subtraction; diameter of largest sphere: 2.60-2.90 µm). To compare CD68+ vesicle properties based on their cellular position, the Iba1 channel/image was “masked” and used to generate a surface to only cover the Iba1 body where thin, processes extend from, defined as the main body, using the Machine Learning Segmentation function. The manual training was iteratively adjusted and then manually reviewed to ensure that the glial body was properly covered and no processes were included. Surfaces were individually and manually edited to connect any segments that were segregated, but were in fact part of the main body, using “unify” function, and likewise, we removed any parts of the surface that covered thin processes extending from the main body using the “cut surface” function. CD68+ surfaces were then parsed into “CD68+ vesicles in the main body” and “CD68+ vesicles in the processes” using the classification feature based on their distance from the generated main body surface. Surfaces inside and/ or in contact with the main body surface (0 µm distance) were classified as “CD68+ vesicles in the main body”, while the remaining vesicles were classified into the latter group. CD68+ vesicle properties including individual vesicle volume (µm³) and CD68 enrichment (mean CD68 intensity/vesicle) as well as the protease enrichment (mean intensity of cathepsin L or B within each individual CD68+ vesicle) were computed using IMARIS.

#### LAMP1+ Analysis

30-μm coronal hemibrain sections from 8-month-old AP-4 WT and KO male mice, obtained as described above, were stained for Iba1, CD68, and cathepsin L. Z-stack images that encompassed the Iba1 signal (typically 200-300 planes with 0.1 µm step size) and were within a similar CA1 region of the hippocampus between AP-4 WT and KO mice, were analyzed using IMARIS 10.2 (Oxford Instruments). Iba1 signal correction, glial cell body isolation using the Machine Learning Segmentation function for surfaces was performed as described above. Once the glial cell bodies were isolated, the LAMP1 channel/image was “masked” within finalized Iba1 surfaces, where signal from only within the completely rendered Iba1 surfaces is used to isolate LAMP1+ vesicles within these whole glia. LAMP1 surfaces were generated in the LAMP1 mask using automated surfaces (background subtraction; diameter of largest sphere: 2.00 µm). Comparison of LAMP1+ vesicle properties based on their cellular position (main body versus processes), was done as described above. LAMP1+ vesicle properties including volume (µm³) and LAMP1 enrichment (mean LAMP1 intensity/vesicle) as well as the protease enrichment (mean intensity of cathepsin L within each individual LAMP1+ vesicle) were computed using IMARIS.

#### Evaluation of number of glial processes

Counting of primary glial processes (the processes that directly emerge from the main body of each microglia) was carried out manually on Iba1-stained coronal brain section images of 8-month-old AP-4 WT and KO male mice using IMARIS software. Utilizing the whole glial and main body surfaces generated from Iba1 staining, which we used in analyzing CD68 vesicle properties, glial processes belonging to each whole/intact glia were identified in the Iba1 mask channel and counted. Both the mean number of processes per glia for an animal and individual glia data were computed.

#### Evaluation of glia size

Glial size (µm³) was analyzed on Iba-1-stained coronal brain section images of 8-month-old AP-4 WT and KO male mice using IMARIS software. Utilizing the Iba1 3-D surface renderings we had generated above, the total volume of each individual glia/surface was computed automatically using IMARIS. We also calculated the mean glial volume for each individual animal.

### Statistical Methods

Graphs were plotted using GraphPad Prism 10 software. Data were represented as Superplots that include mean ± SEM as well as individual data points unless otherwise specified. GraphPad Prism 10 was used to perform unpaired t-test for figures 1B, 2C-E, 3B, F, I and J as well as supplementary figures 2D and 3E. The different shapes (circles, squares, triangles, and diamonds), each represented an individual animal. For all other experiments, we fitted a linear mixed effects (LME) model (Smith et al., 2026; Spivey et al., 2026) and the statistical significance was determined by Wald t-test. Our data in these instances, have a nested structure, with multiple individual vesicles or cells (technical replicates) measured within each biological replicate (brain tissue from different individual mice) that have one of the following fixed effects: Cellular Position /location within glia (Fig. 1C, D, F, Fig. S1 B, Fig. 2H, K, N, Fig. S2 I, Fig. 3C, D, G, H, Fig. S3 B, D), or Genotype (Fig. 2F, G, I, J, L, M, Fig. S2 E-H, Fig. 3K-N, Fig. S3 F, G). To account for the non-independence between the technical replicates within each biological replicate, we fit a linear mixed-effects (LME) model with biological replicates defined as a random effect. This model allows us to assess genotype/cellular location-level effects while preserving the variability of individual vesicles or cells within the hierarchical structure of the data. We used the RStudio (Version 2025.05.1+513) for performing the LME model (LME, R package “nlme”) or LME model with “weights” argument (R package “nlme”, weight = varIdent). Model residuals were inspected for homoscedasticity. For heteroscedastic data, model fit was improved by incorporating a varIdent variance structure. Figures that have heteroscedastic data include figures 1C, D, F, 2F-N, 3C, D, G, H, K-N, and supplementary figures 1B, 2G-I, 3B, D, F, and G. Figures with homoscedastic data are supplementary figures 2E and F. Use of AI ChatGPT was used to aid writing the code in R for the statistical analysis. The authors carefully reviewed and edited all AI-generated edits and take full responsibility for the content in the publication.

## Acknowledgements

We thank Jose Saltos and Rameen Tahir for help with mouse colony maintenance. Grants from the NIH(RF1AG076653,R01AG076653,R01AG074248) to S.G. provided financial support for the lab’s research focused on lysosome biology. E.S. was a recipient of a T32 fellowship, from the Training program in the biology and translational research on Alzheimer’s disease and related dementias (T32AG057468) from July 2024-August 2026. We also thank the IMARIS support team for their input on image analysis using IMARIS software. Figures that were created in BioRender: Created in BioRender. Somodji, E. (2026) https://core.local.biorender.dev/api/short-link/ptvjpz7.

## Notes

### Competing Interest Statement

The authors have declared no competing interest.

