## Supplemental Figs and Legends for "Compartment-Specific Lysosomal Heterogeneity in Microglia Is Regulated by Adaptor Protein Complex-4"

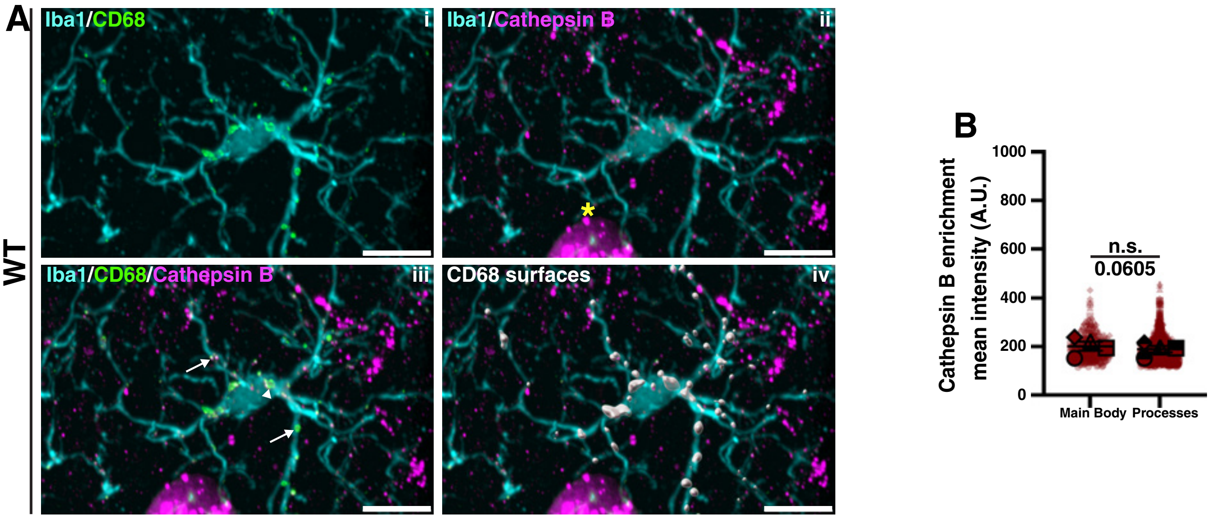
A
WT
Iba1/CD68
i
Iba1/Cathepsin B
ii
*
Iba1/CD68/Cathepsin B
iii
CD68 surfaces
iv
B
Cathepsin B enrichment mean intensity (A.U.)
1000
800
600
400
200
0
n.s.
0.0605
Main Body
Processes

**Figure S1**

**Figure S1. CD68+ vesicles in WT microglia in distinct subcellular positions, do not differ in cathepsin B enrichment**

(A) High-resolution images highlighting (i) a microglia (Iba1; cyan) in the WT hippocampus CA1 region co-stained with CD68 (green) labeling endo-lysosomes within the glia; (ii) image highlighting the cathepsin B (magenta), present within the glia (Iba 1; cyan) and also in neighboring neurons (yellow asterisk). (iii) Merged image of Iba1, CD68 and cathepsin B highlighting vesicles within the main body (white arrowheads) and processes (white arrows) of glia. (iv) Image of Iba1, CD68 and cathepsin B, where the isolated CD68 vesicles from within the individual glia (identified by its Iba1 mask) are rendered in 3D or as white "surfaces". The cathepsin B content from within these surfaces are specifically analyzed to capture properties relating to the individual microglia's vesicles. Bar, 10 µm. (B) Superplot depicting cathepsin B enrichment, as measured by mean intensity of cathepsin B (arbitrary intensity units), of all individual CD68+ vesicles from the main body and processes of microglia of the individual WT animals (circle, square, triangle, and diamond), as well as the mean cathepsin B intensity of each animal (larger symbol with bold outline) ± SEM. N = 4 animals, n = 341 CD68+ vesicles in the main body and n = 1,491 CD68+ vesicles in the processes. ns = not significant. Data was analyzed using an LME model.

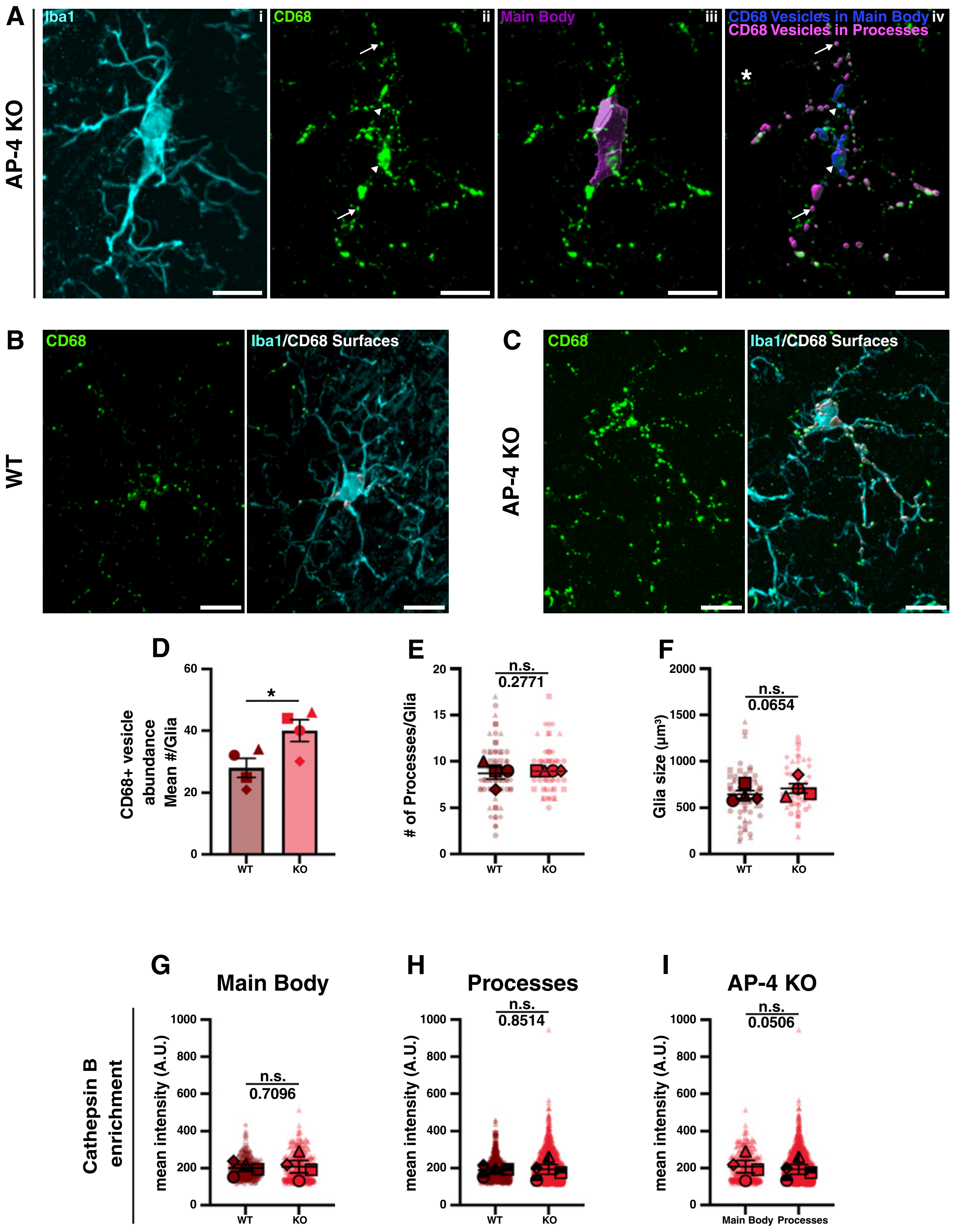
A
AP-4 KO
Iba1
i
CD68
ii
Main Body
iii
CD68 Vesicles in Main Body
iv
CD68 Vesicles in Processes
B
WT
CD68
Iba1/CD68 Surfaces
C
AP-4 KO
CD68
Iba1/CD68 Surfaces
D
CD68+ vesicle abundance Mean #/Glia
60
40
20
0
*
WT
KO
E
### of Processes/Glia
20
15
10
5
0
n.s.
0.2771
WT
KO
F
Glia size (µm³)
2000
1500
1000
500
0
n.s.
0.0654
WT
KO
G
Main Body
Cathepsin B enrichment
mean intensity (A.U.)
1000
800
600
400
200
0
n.s.
0.7096
WT
KO
H
Processes
mean intensity (A.U.)
n.s.
0.8514
WT
KO
I
AP-4 KO
mean intensity (A.U.)
n.s.
0.0506
Main Body
Processes

Figure S2

**Figure S2. Loss of AP-4 function does not alter glia size, number of glial processes, nor cathepsin B content within CD68+ vesicles**

(A) Representative high-resolution images depicting a single AP-4 KO microglia. Iba1 [(i) glial body; cyan] was used to identify microglial cell bodies, while CD68 [(ii) CD68+ vesicles; green] was used to identify microglial endo-lysosomes in the main body (white arrowheads) and processes (white arrows). (iii) Image showing “main body” (purple surface) of microglia and CD68+ vesicles (green). (iv) Image showing CD68+ vesicles in 3D or as rendered “surfaces” within the main body (arrowheads; blue surfaces) and in processes (arrows; magenta surfaces). Note, CD68+ vesicles outside the individual microglia’s body are in green (white asterisk) and not included in the glia’s CD68+ vesicle pool. Bar, 10 µm. (B, C) Representative high-resolution images of microglia in hippocampus CA1 region of AP-4 WT (B) and KO mice (C) stained for microglial cell bodies (Iba1; cyan) and microglial vesicles (CD68; green). Isolated CD68+ vesicles from within the glia (identified by its Iba1 mask) are depicted as white surfaces. Bar, 10 µm. (D) Quantification of mean number of CD68+ vesicles per microglia within whole glia of WT and KO glia in the hippocampus CA1 region of 8-month-old male AP-4 WT and KO animals (WT – brown; KO – red). Each shape (circle, square, triangle, and diamond) represents data from an individual animal. Mean ± SEM, N = 4 animals, n = 131 WT glia, n = 109 AP-4 KO glia. * $p<0.05$, unpaired *t*-test**.** (E) Superplot depicting the number of primary glial processes per individual microglia from the individual AP-4 WT and KO animals (circle, square, triangle, and diamond), as well as the mean number of primary glial processes of each animal (larger symbol with bold outline) ± SEM. N = 4 animals, n = 66 WT glia, n = 57 AP-4 KO glia. ns = not significant. Data was analyzed using an LME model. (F) Superplot depicting the glia size, as measured by volume ($\mu m^3$), of all individual microglia from individual AP-4 WT and KO animals (circle, square, triangle, and diamond), as well as the mean glial volume of each animal (larger symbol with bold outline) ± SEM. N = 4 animals, n = 66 WT glia, n = 57 AP-4 KO glia. ns = not significant. Data was analyzed using an LME model. (G) Superplot depicting cathepsin B enrichment, as measured by mean intensity of cathepsin B (arbitrary intensity units), of all individual CD68+ vesicles from the main bodies of microglia from individual AP-4 WT and KO animals (circle, square, triangle, and diamond), as well as the mean cathepsin B intensity of each animal (larger symbol with bold outline) ± SEM. N = 4 animals, n = 341 CD68+ vesicles in WT main bodies and n = 330 CD68+ vesicles in AP-4 KO main bodies. ns = not significant. Data was analyzed using an LME model. (H) Superplot depicting cathepsin B enrichment, as measured by mean intensity of cathepsin B (arbitrary intensity units), of all individual CD68+ vesicles from the processes of microglia from individual AP-4 WT and KO animals (circle, square, triangle, and diamond), as well as the mean cathepsin B intensity of each animal (larger symbol with bold outline) ± SEM. N = 4 animals, n = 1,491 CD68+ vesicles in WT processes and n = 1,743 CD68+ vesicles in AP-4 KO processes. ns = not significant. Data was analyzed using an LME model. (I) Superplot depicting cathepsin B enrichment, as measured by mean intensity of cathepsin B (arbitrary intensity units), of all individual CD68+ vesicles from the main body and processes of microglia from individual AP-4 KO animals (circle, square, triangle, and diamond), as well as the mean cathepsin B intensity of each animal (larger symbol with bold outline) ± SEM. N = 4 animals, n = 330 CD68+ vesicles in the main body and n = 1,743 CD68+ vesicles in the processes. ns = not significant. Data was analyzed using an LME model.

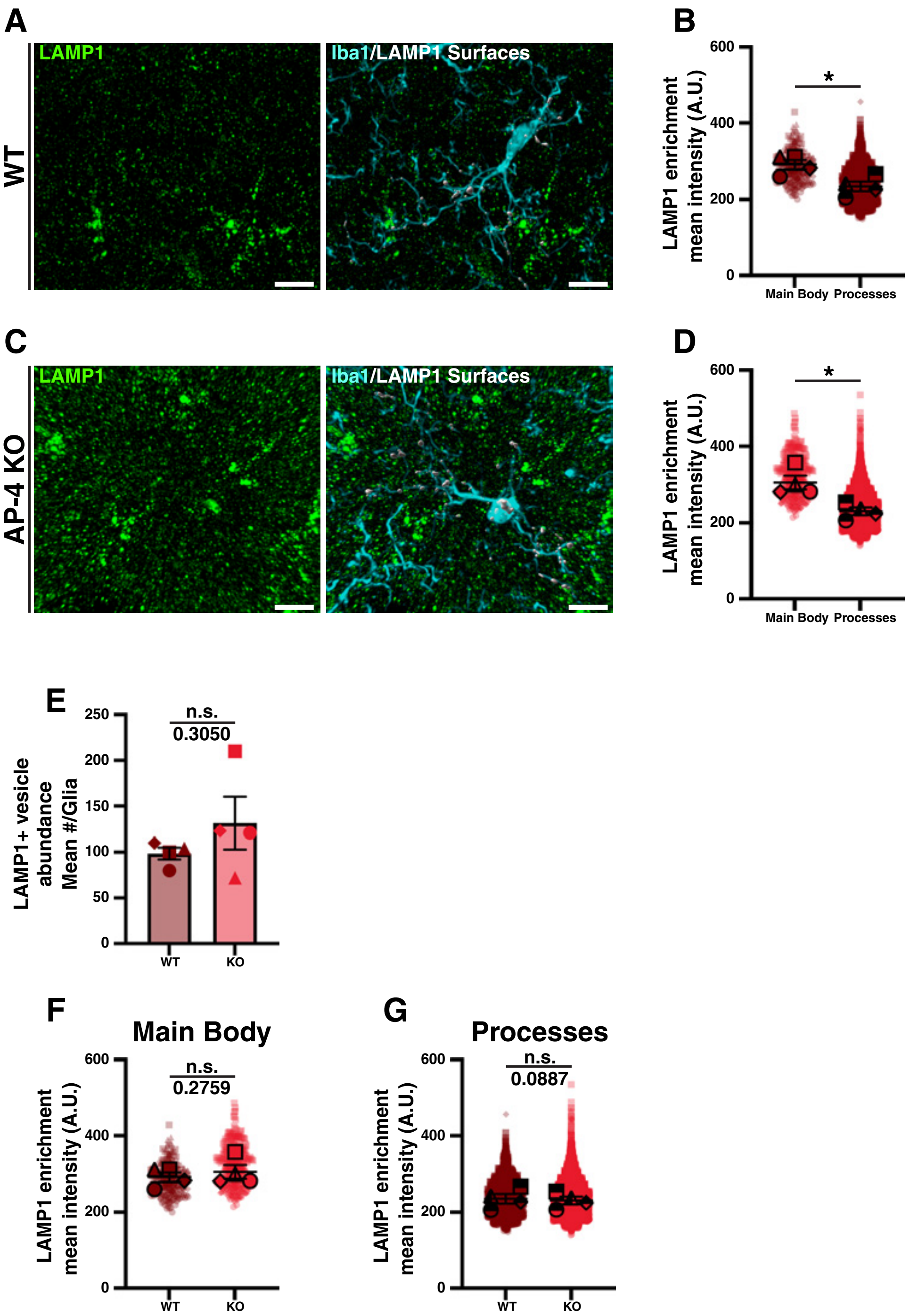
A
WT
AP-4 KO
LAMP1
Iba1/LAMP1 Surfaces
B
600
400
200
0
*
LAMP1 enrichment mean intensity (A.U.)
Main Body
Processes
C
D
E
LAMP1+ vesicle abundance Mean #/Glia
250
150
100
50
n.s.
0.3050
WT
KO
F
Main Body
0.2759
G
Processes
0.0887

Figure S3

**Figure S3. Loss of AP-4 does not impact LAMP1+ vesicle abundance or LAMP1 enrichment**

(A) Representative high-resolution image of a microglia from hippocampus CA1 region of an AP-4 WT mouse stained to label the whole microglial body (Iba1; cyan) and endo-lysosomes (LAMP1; green) present both within and outside the glia. Use of the rendered Iba1 surface as a mask helps isolate LAMP1+ vesicles from within the individual glia (white surfaces). Bar, 10 µm. (B) Superplot depicting LAMP1 enrichment, as measured by mean intensity (arbitrary intensity units), of all individual LAMP1+ vesicles from the main body and processes of microglia from individual WT animals (circle, square, triangle, and diamond), as well as the mean LAMP1 intensity of each animal (larger symbol with bold outline) ± SEM. N = 4 animals, n = 246 LAMP1+ vesicles in the main body; n = 3,727 LAMP1+ vesicles in the processes. * $p<0.05$. Shown are *p*-values from an LME model. (C) Representative high-resolution image of an AP-4 KO microglia from hippocampus CA1 region of an AP-4 KO mouse stained for the microglial cell body (Iba1; cyan) and endo-lysosomes (LAMP1; green) present within and outside the glia. Use of the rendered Iba1 surface as a mask helps isolate LAMP1+ vesicles from within the individual glia (white surfaces). Bar, 10 µm. (D) Superplot depicting LAMP1 enrichment, as measured by mean intensity (arbitrary intensity units), of all individual LAMP1+ vesicles from the main body and processes of microglia from individual AP-4 KO animals (circle, square, triangle, and diamond), as well as the mean LAMP1 intensity of each animal (larger symbol with bold outline) ± SEM. N = 4 animals, n = 423 LAMP1+ vesicles in the main body; n = 7,625 LAMP1+ vesicles in the processes. * $p<0.05$. Shown are *p*-values from an LME model. (E) Quantification of mean number of LAMP1+ vesicles per microglia within whole WT and KO glia in the hippocampus CA1 region of 8-month-old male AP-4 WT and KO animals (WT – brown; KO – red). Each shape (circle, square, triangle, and diamond) represents data from an individual animal. Mean ± SEM, N = 4 animals, n = 42 WT glia, n = 57 AP-4 KO glia. ns = not significant, unpaired *t*-test**.** (F) Superplot depicting LAMP1 enrichment, as measured by mean intensity (arbitrary intensity units), of all individual LAMP1+ vesicles from the main bodies from microglia of the individual AP-4 WT and KO animals (circle, square, triangle, and diamond), as well as the mean LAMP1 intensity of each animal (larger symbol with bold outline) ± SEM. N = 4 animals, n = 246 LAMP1+ vesicles in WT main bodies; n = 423 LAMP1+ vesicles in AP-

4 KO main bodies. ns = not significant. Data was analyzed using an LME model. (G) Superplot depicting LAMP1 enrichment, as measured by mean intensity (arbitrary intensity units), of all individual LAMP1+ vesicles from the processes from microglia of the individual AP-4 WT and KO animals (circle, square, triangle, and diamond), as well as the mean LAMP1 intensity of each animal (larger symbol with bold outline) ± SEM. N = 4 animals, n = 3,727 LAMP1+ vesicles in WT processes; n = 7,625 LAMP1+ vesicles in AP-4 KO processes. ns = not significant. Data was analyzed using an LME model.

**Supplement Video Legends**

**Video 1. Isolation of microglia using IMARIS surface feature**

IMARIS-generated video demonstrating the rendering of Iba1 surfaces to isolate individual microglia (Iba1; cyan) from high-resolution z-stack images of the hippocampus CA1 region of the mouse brain, using IMARIS “surfaces” feature. Bar, 20 µm.

**Video 2. Isolation of microglia and CD68 vesicles within them using IMARIS surfaces**

IMARIS-generated video demonstrating isolation of microglia (Iba1; cyan) in the hippocampus CA1 region of the mouse brain using surfaces. Use of rendered surfaces that are complete in the z-stack as a “mask” helps display only the isolated glia and their CD68+ vesicles (green) which are in turn rendered as individual vesicle “surfaces”, seen in white.

**Video 3. Identification and characterization of CD68+ vesicles based on subcellular position in WT microglia through IMARIS**

IMARIS-generated video showing an example of how CD68+ vesicles in main body (blue) and CD68+ vesicles in processes (magenta), are parsed from CD68+ vesicles (green) within WT microglia, once their main body is identified. Bar, 10 µm.

**Video 4. Evaluation of cathepsin enrichment/content specifically from microglial CD68+ vesicles**

IMARIS-generated video showing how cathepsin B content (magenta) from only glial CD68+ vesicles (green) can be isolated once the CD68+ surfaces (white) surfaces are rendered within the individual glia (Iba1; cyan). Bar, 10 µm.

**Video 5. Identification and characterization of CD68+ vesicles based on subcellular position in AP-4 KO microglia through IMARIS**

IMARIS-generated video showing an example of how CD68+ vesicles in main body (blue) and CD68+ vesicles in processes (magenta), are parsed from CD68+ vesicles (green) within AP-4 KO microglia, once their main body is identified. Bar, 10 µm.

**Video 6. 3D view of a representative AP-4 WT microglia stained for Iba1 and CD68**

IMARIS-generated video showing a 3D, 360-degree view of a representative masked WT microglia from the hippocampus CA1 region (Iba1; cyan) and its individual CD68+ vesicles (CD68; green). CD68+ vesicles isolated using surfaces (white). Bar, 10 µm.

**Video 7. 3D view of a representative AP-4 KO microglia stained for Iba1 and CD68**

IMARIS-generated video showing a 3D, 360-degree view of a representative masked AP-4 KO microglia within the hippocampus CA1 region (Iba1; cyan) and its individual CD68+ vesicles (CD68; green). CD68+ vesicles isolated using surfaces (white). Bar, 10 µm.

**Video 8. 3D view of a representative AP-4 WT microglia stained for Iba1 and LAMP1**

IMARIS-generated video showing a 3D, 360-degree view of a representative masked WT microglia within the hippocampus CA1 region (Iba1; cyan) and its individual LAMP1+ vesicles (LAMP1; green). LAMP1+ vesicles isolated using surfaces (white). Bar, 10 µm.

**Video 9. 3D view of a representative AP-4 KO microglia stained for Iba1 and LAMP1**

IMARIS-generated video showing a 3D, 360-degree view of a representative masked AP-4 KO microglia within the hippocampus CA1 region (Iba1; cyan) and its individual LAMP1+ vesicles (LAMP1; green). LAMP1+ vesicles isolated using surfaces (white). Bar, 10 µm.
